# morse: an R-package in support of Environmental Risk Assessment

**DOI:** 10.1101/2021.04.07.438826

**Authors:** Virgile Baudrot, Sandrine Charles

## Abstract

Package `morse` is devoted to the analysis of experimental data collected from standard toxicity tests. It provides ready-to-use functions to visualize a data set and to estimate several toxicity indices to be further used in support of environmental risk assessment in full compliance with regulatory requirements. Such toxicity indices are indeed classical requested by standardized regulatory guidelines on which national agencies base their evaluation of applications for marketing authorisation of chemical active substances.

Package `morse` can be used to get estimates of *LC_x_* (*x*% Lethal Concentration) or *EC_x_* (*x*% Effective Concentration) by fitting standard exposure-response models on toxicity test data. Risk indicator estimates as well as model parameters are provided along with the quantification of their uncertainty. Package `morse` can also be used to get estimates of the *NEC* (No Effect Concentration) by fitting a Toxicokinetic-Toxicodynamic (TKTD) model (namely `GUTS` models, that is *General Unified Threshold models of Survival*). Using `GUTS` models also allow to get estimates of *LC*_(*x,t*)_ (whatever *x* and *t*) and *LP*_(*x,t*)_, this later being defined by EFSA as the x% multiplication factor leading to an additional reduction of x% in survival at the end of the exposure profile. Above all, `GUTS` models can be used on data collected under time-variable exposure profiles.

This paper illustrates a typical use of `morse` with survival data collected over time and at different increasing exposure concentrations, analysed with the reduced version of GUTS models based on the stochastic death hypothesis (namely, the `GUTS-RED-SD` model). This example can be followed step-by-step to analyse any new data set, as long as the data set format is respected.

## Statement of Need

Package `morse` (Baudrot et al., 2021) has been tested using R (version 3.5 and later) on macOS, Linux and Windows machines. Regarding the particular case of toxicokinetic-toxicodynamic (TKTD) models for survival, namely GUTS models (Jager & Ashauer, 2018), package `morse` was ring-tested together with nine other `GUTS` implementations under different software platforms. All participants to the ring-test received the same data sets and tasks, carried out their simulations independently from each other and sent the results back to the coordinator for analysis. Giving very similar results than the other implementations, package `morse` was thus confirmed as fit-for-purpose in fitting GUTS models on survival toxicity test data.

All functions in package `morse` can be used without a deep knowledge of their underlying probabilistic model or inference methods. Rather, they were designed to behave as well as possible, without requiring the user to provide values for some obscure parameters. Nevertheless, models implemented in `morse` can also be used as a first step to tailor new models for more specific situations.

**Figure 1:**
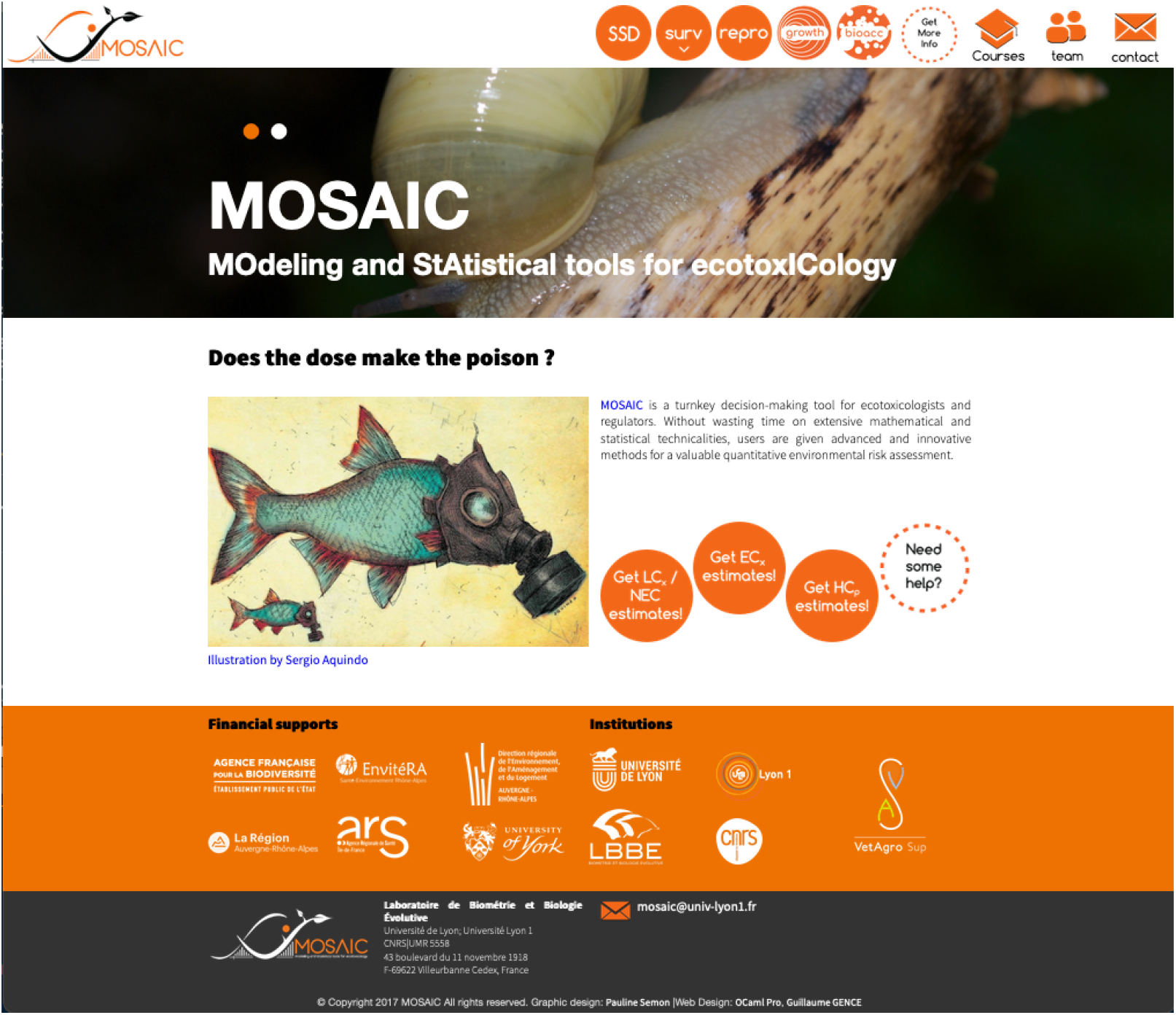
Homepage of the MOSAIC web platform (https://mosaic.univ-lyon1.fr/).

Note that package `morse` benefits from a web interface, MOSAIC, from which the same analyses can be reproduced directly on-line without needs to invest in `R` programming. MOSAIC is freely available at https://mosaic.univ-lyon1.fr/ (Charles, Delignette-Muller, Veber, & Delignette-Muller, 2018) (Figure **??**).

## Availability

Package `morse` is available as an `R` package; it can be directly downloaded from CRAN https://CRAN.R-project.org/package=morse, where package dependencies and system requirements are also documented. The development version can be found on GitHub https://github.com/pveber/morse, where code contributions, bug reports, fixes and feature requests are more than welcome by opening issues and pull requests.

## Main features

The main functions in package `morse` are `survData()`, `reproData()` and `plotDoseResponse()` to visualize raw data. Functions `survFitTT()`, `reproFitTT()`, `survFit()` allow to fit a model on data in order to estimate toxicity indicators, the choice depending on the type of data. Fitting outputs can be either displayed with `plot()` or synthesized with `summary()`. Functions are available to check the goodness-of-fit, namely `ppc()` and `plot_prior_post()`. Predictions can be performed with `predict()`, `predict_ode()`, `predict_Nsurv()` and `predict_Nsurv_ode()`. At last, function `LCx()` and `MFx()` allow to get *x*% lethal concentrations or profiles, respectively.

The `morse` package currently handles binary and count data, as for example survival and reproduction data. Functions dedicated to binary (resp. count) data analysis start with a `surv` (resp. `repro`) prefix. `morse` provides a similar workflow in both cases:

1. create and validate a data set;
2. explore a data set;
3. plot a data set;
4. fit a model on data and get parameter estimates;
5. check goodness-of-fit with Posterior Predictive Check plot (PPC).

In addition, for binary data handled with `GUTS` models, package `morse` also allows to:

1. calculate and plot *LC*_(*x, t*)_ and *LP*_(*x, t*)_;
2. compute goodness-of-fit criteria: the PPC percentage, the Normalized Root Mean Square Error (NRMSE) and the Survival probability prediction error at the end of the exposure profile (SPPE).

See (EFSA PPR Panel, 2018) for details.

Those steps are presented in depth in the `Tutorial` available at https://cran.r-project.org/web/packages/morse/vignettes/tutorial.html, with all necessary details to plenty use all `morse` features. A more formal description of the models and the estimation procedures are provided in a document called “Models in `morse` package” available at https://cran.r-project.org/web/packages/morse/vignettes/modelling.pdf. Please refer to this documentation for further introduction to the use of the `morse` package.

## Minimal Working Example

### Loading morse and its dependencies

In order to use package `morse`, you need to install it with all its dependencies, including JAGS and C++ (see below), as well as other R-packages: mandatory ones (`coda`, `deSolve`, `dplyr`, `epitools`, `graphics`, `grDevices`, `ggplot2` (⩾ 2.1.0), `grid`, `gridExtra`, `magrittr`, `methods`, `reshape2`, `rjags` (⩾ 4.0), `stats`, `tibble`, `tidyr`, `zoo`) and suggested ones (`knitr`, `rmarkdown`, `testthat`). For this purpose, you can use the two classical R commands:

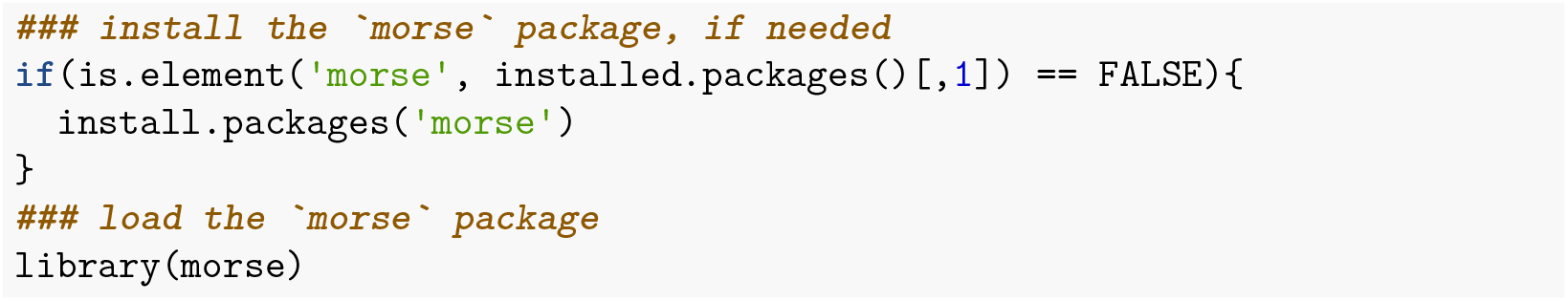

### JAGS

The `morse` package is linked to JAGS http://mcmc-jags.sourceforge.net/ that is the Bayesian sampler used to perform inference with all implemented models. So, you need also to download and install JAGS at https://sourceforge.net/projects/mcmc-jags/. Then you must test that your R graphical user interface has access to JAGS, and, if not, to specify where JAGS can be found on your computer. Indeed, once installed, JAGS can be lost in the PATH. To help solving this issue, you can use package `runjags` which is not within `morse` so that you have to install and load it too.

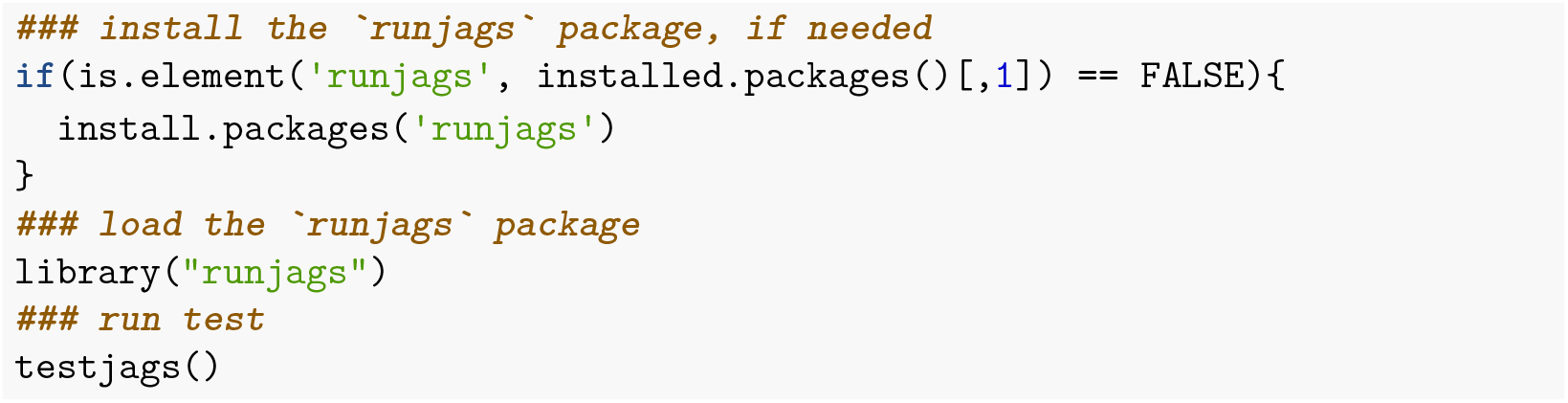

The output should look like this:

~~~
You are using R version 4.0.2 (2020-06-22) on a windows machine, with the RStudio
JAGS version 4.3.0 found successfully using the command
‘C:/Program Files/JAGS/JAGS-4.3.0/x64/bin/jags-terminal.exe’
The rjags package is installed
~~~

Otherwise, you can specify to your R graphical user interface where JAGS executable is located in your computer (somewhere in `‘C:/Program Files/JAGS/JAGS-4.3.0/x64/bin/jags-terminal.exe’` on Windows machines):

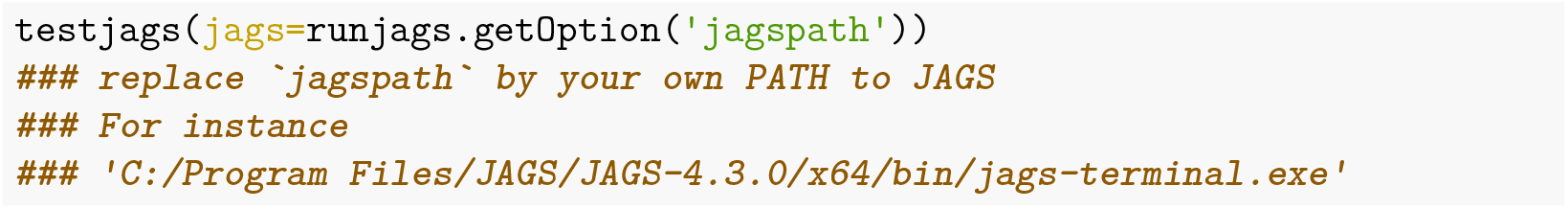

### C++

The `morse` package is also linked to C++. C++ is used for running simulations leading to predictions. In R, you should not have issues with C++ requirements since it is very well integrated (many R functions are simple interfaces to C++ functions). Feel free to report any trouble at https://github.com/pveber/morse/issues by opening a new issue for the `morse` package.

### Survival analysis

We assume hereafter that `morse` and all the above dependencies have been corrected installed. To illustrate the use of `morse`, we will use a standard survival data set coming from a chronic laboratory toxicity test with *Gammarus pulex*, a freshwater invertebrate, exposed to increasing concentrations of propiconazole (a fungicide) during four days. Eight concentrations were tested with two replicates of 10 organisms per concentration. Survival was monitored at five time points (at day 0, 1, 2, 3 and 4) (Nyman, Schirmer, & Ashauer, 2012).

We will used the reduced version of the GUTS model based on the stochastic death hypothesis (namely, the `GUTS-RED-SD` model), as recommended by the *European Food Safety Authority* (EFSA) for the environmental risk assessment (ERA) of plant protection products potentially toxic for aquatic living organisms (EFSA PPR Panel, 2018). This model can also be fitted on-line with the MOSAIC web platform (Baudrot, Veber, Gence, & Charles, 2018). Below is the *modus operandi* with package `morse` to be followed step-by-step in order to be in full compliance with the EFSA workflow for ERA.

#### Calibration step

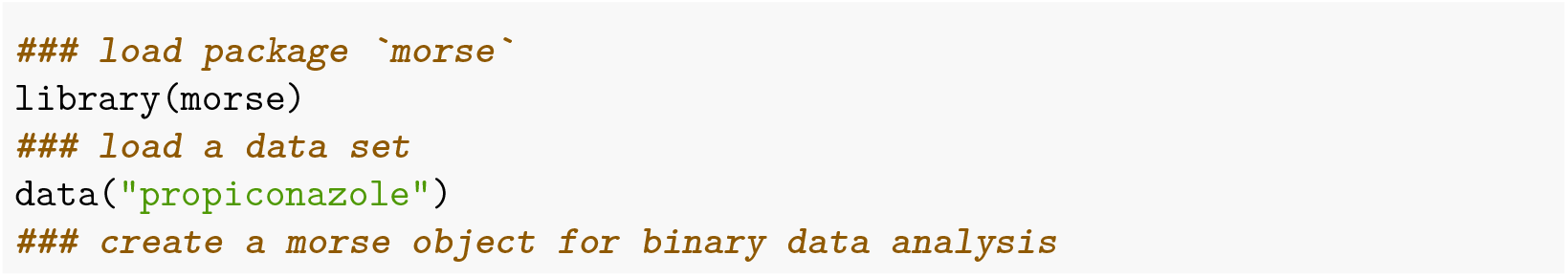

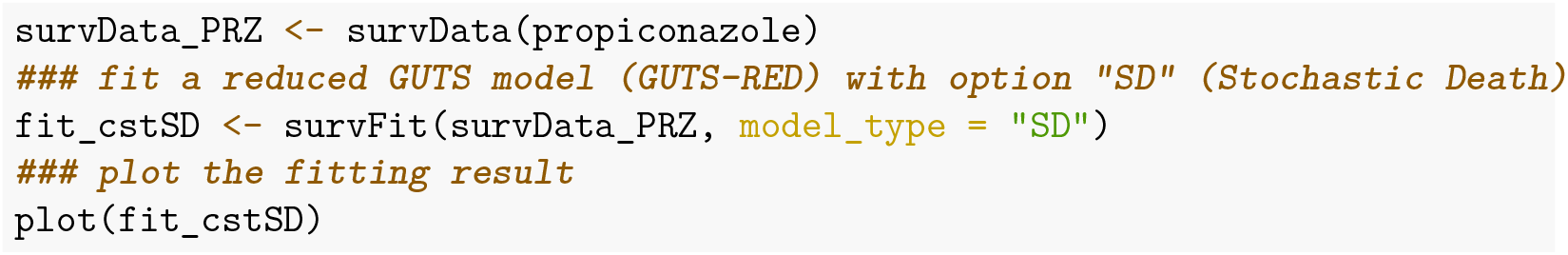

**Figure 2:**
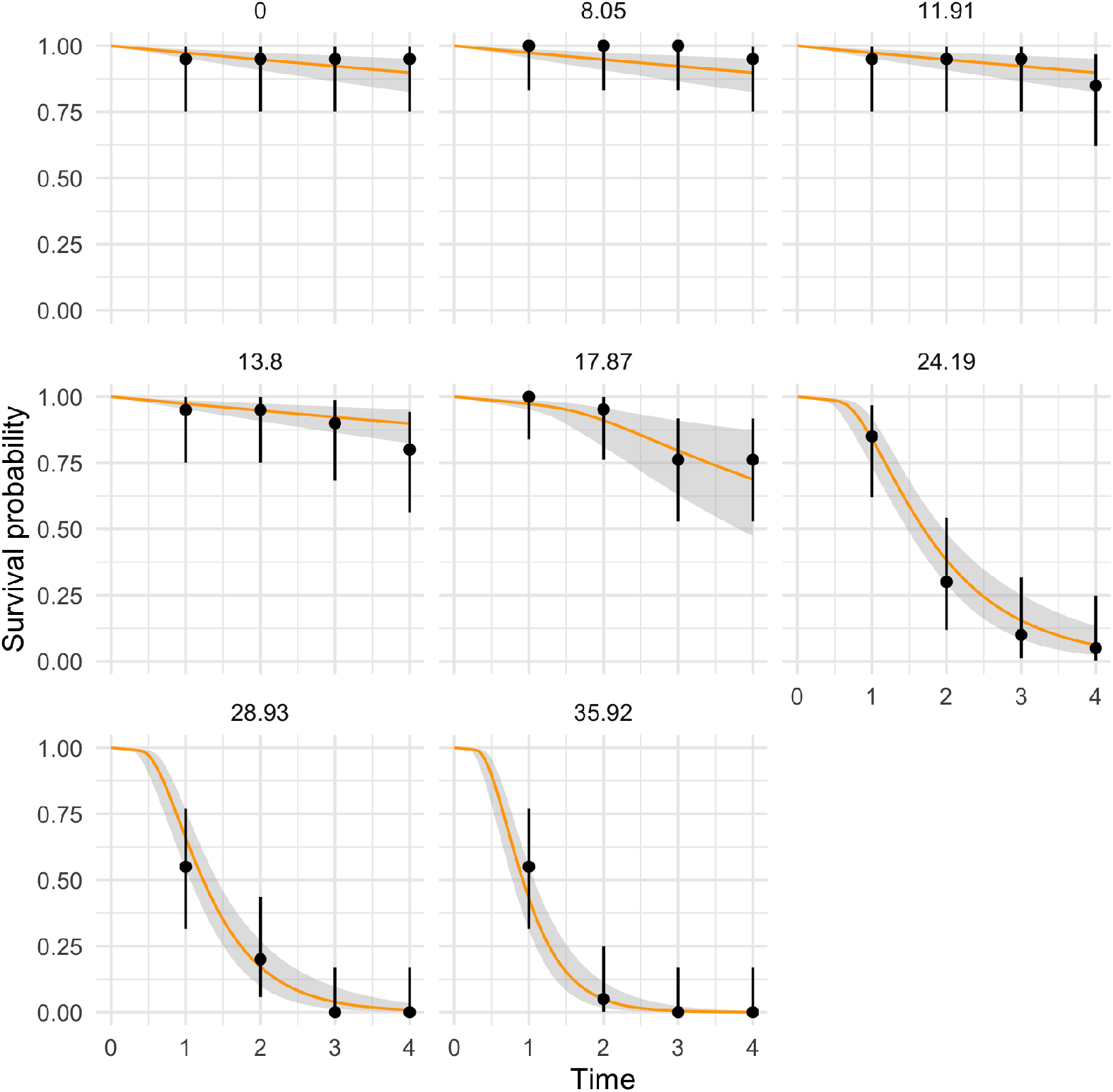
Fitting result with a GUTS-RED-SD model. The median fitted curves are in orange and the uncertainty bands in gray. Black dots are observed data surrounded by their binomial confidence intervals.

#### Get the *x*% lethal concentration

Using a GUTS model with `morse` allows to get a probability distribution on the *x*% lethal concentration whatever the exposure duration *t*, namely the *LC*_(*x,t*)_. By default, *t* corresponds to the last time point in the data set and *x* = 50%.

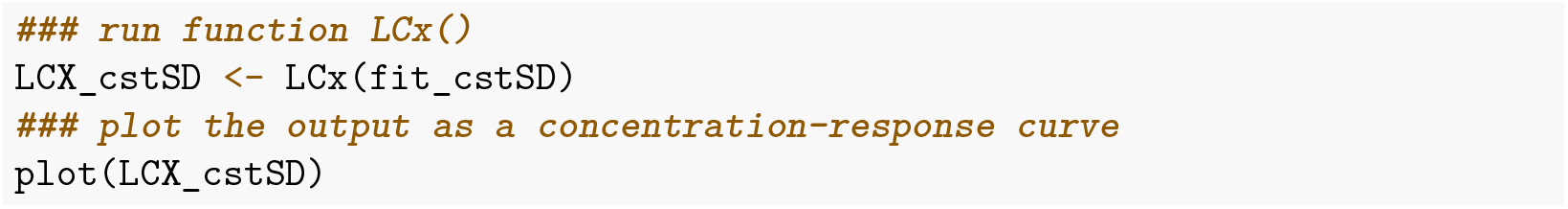

**Figure 3:**
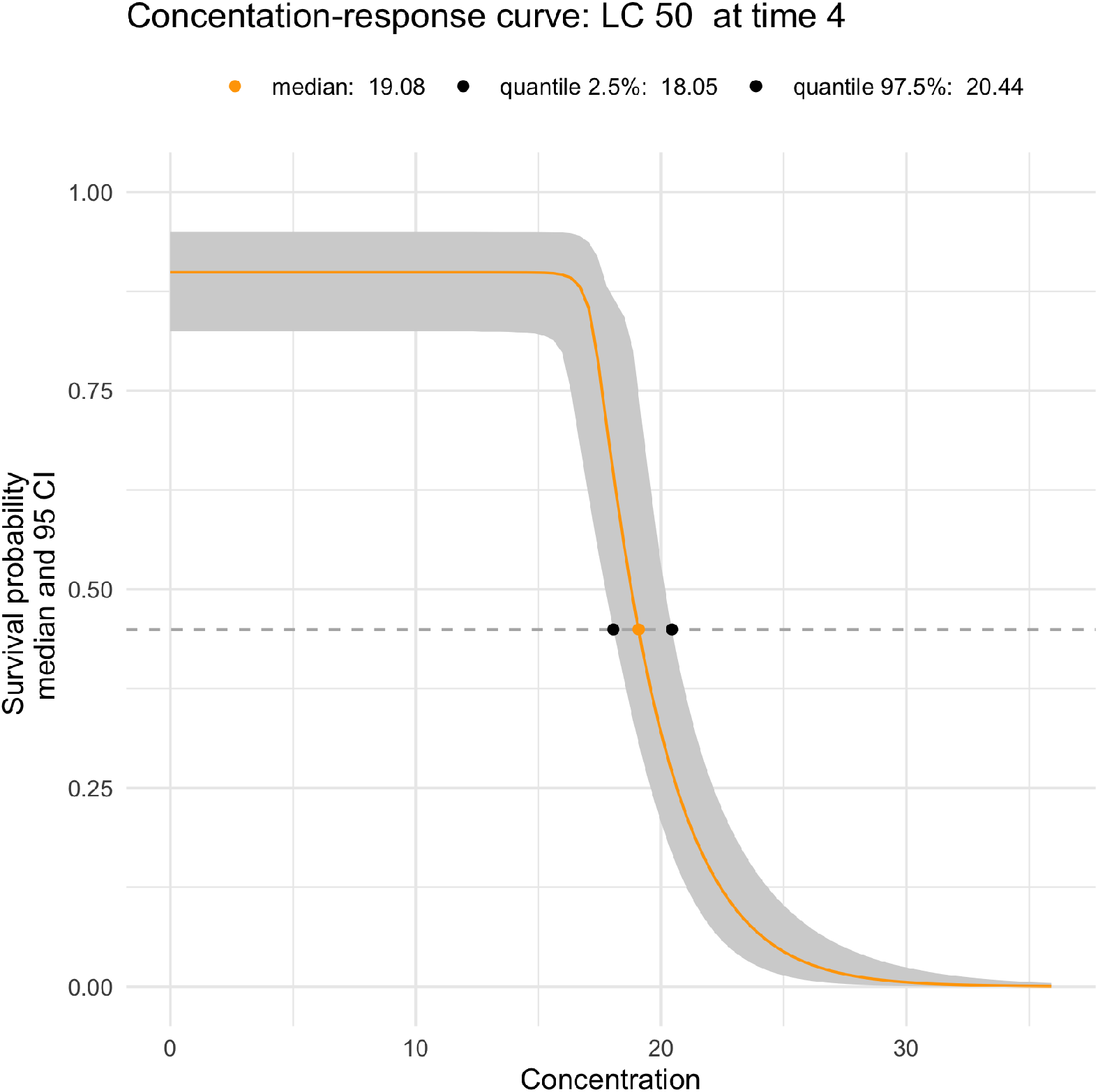
Simulated concentration-response curve corresponding to the previous fitting result on Figure 2.

#### Validation step

Validation consists in predicting the number of survivors over time under pulsed-exposure profiles for which observations have also been collected. Predictions are then compared to observations and their adequacy is checked according to several validation criteria defined by EFSA (EFSA PPR Panel, 2018). The aim of this step is to choose an appropriate model for the following step.

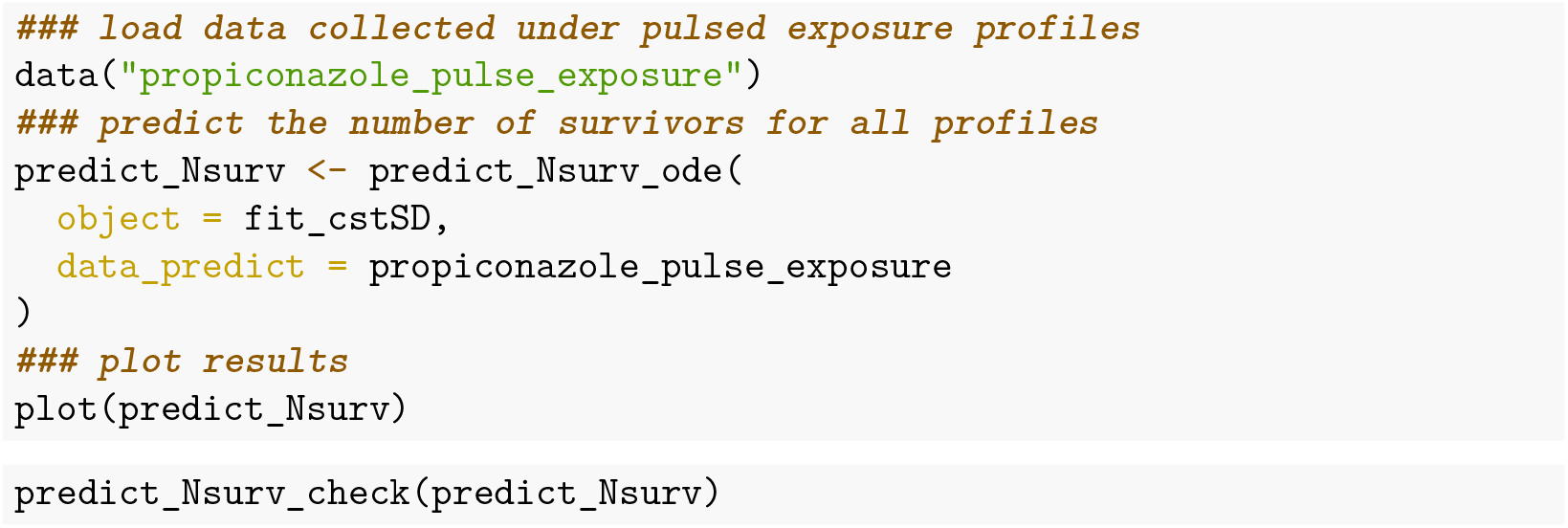

Once the predictions are visually checked (Figure **??**), quantitative validation criteria need to be calculated.

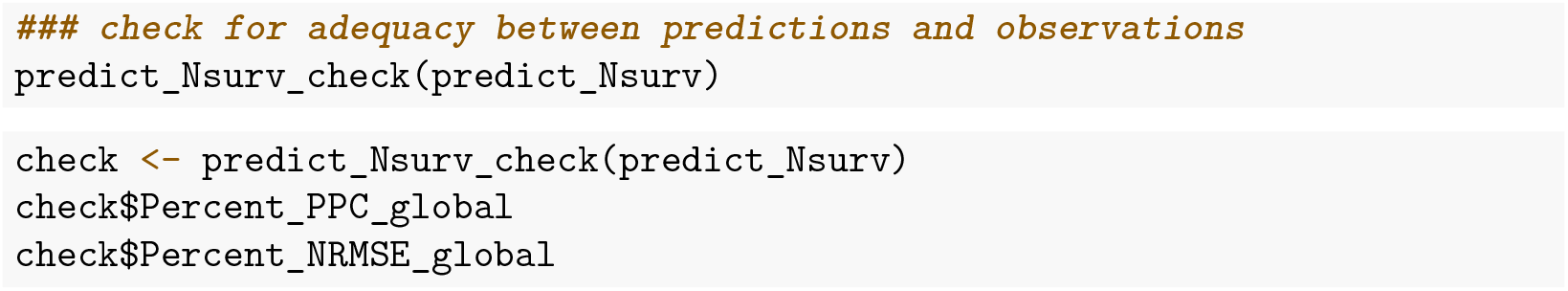

This reveals that, in total, 84% of the observations lie within the uncertainty band of the predictions, while the global variability of data around the predictions is 16.2%. For both criteria, a maximum value of 50% is expected, what means here that we do not expect specific risk for the species and the chemical compound under consideration.

#### Prediction step

Risk assessors are interested in testing various exposure scenarios, having a certain environmental realism that is varying over time. Risk assessors expect to evaluate the potential impact of these profiles on survival of target species to protect. Typically, they want to compute the multiplication factor *MF*_(*x, t*)_ that could be applied to the exposure profile without reducing more than by *x*% the survival probability at a specified test duration t (default being the last time point of the exposure profile). This is the so-called *x*% lethal profile, denoted *LP_x_*, and newly proposed by (EFSA PPR Panel, 2018). This calculation is provided by function `MFx()` in `morse`.

The mathematical definition of the *x*% Multiplication Factor at time *t* (at the end of a time series *T* = {0, *t*}) is given by:

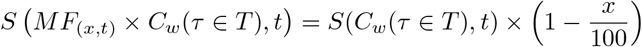

where *C_w_*(*τ* ∈ *T*) is the original exposure profile, and expression *S*(*MF*_(*x,t*)_ × *C_w_*(*τ* ∈ *T*), *t*) the survival probability after the exposure profile has been translated upward by a multiplication *MF*_(*x,t*)_; the new exposure profile thus becomes equal to *MF*_(*x,t*)_ × *C_w_*(*τ* ∈ *T*).

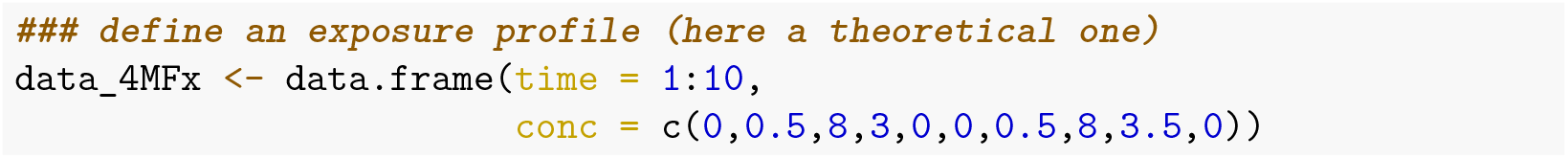

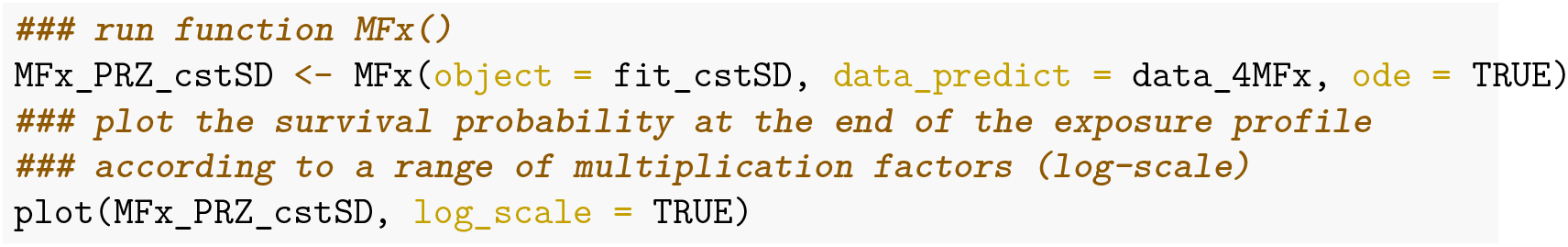

**Figure 4:**
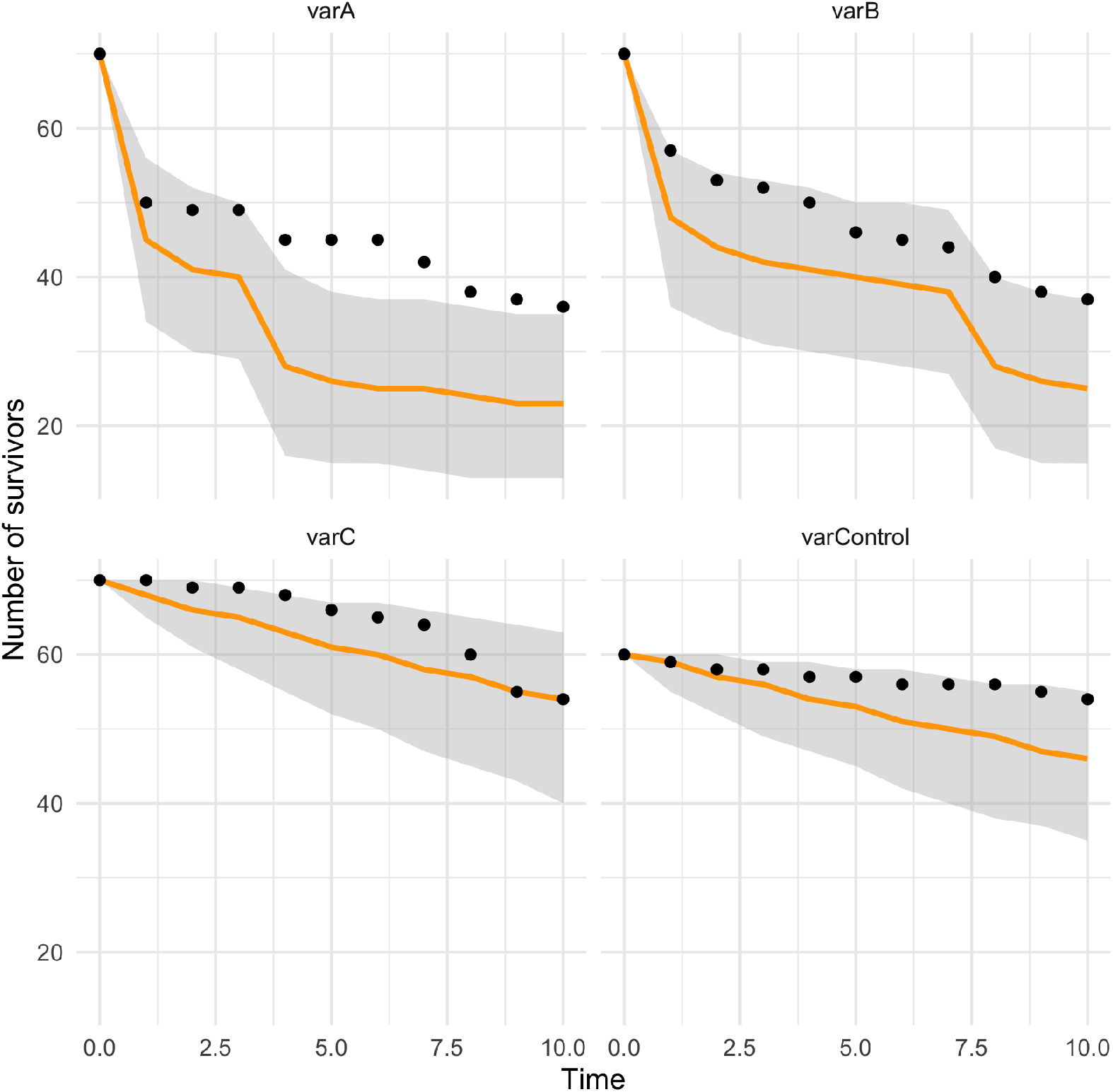
Visual check of adequacy between predictions based on the GUTS-RED-SD model with parameter values estimated in the calibration step (median prediction in orange, uncertainty band in gray), and observations (black dots).

**Figure 5:**
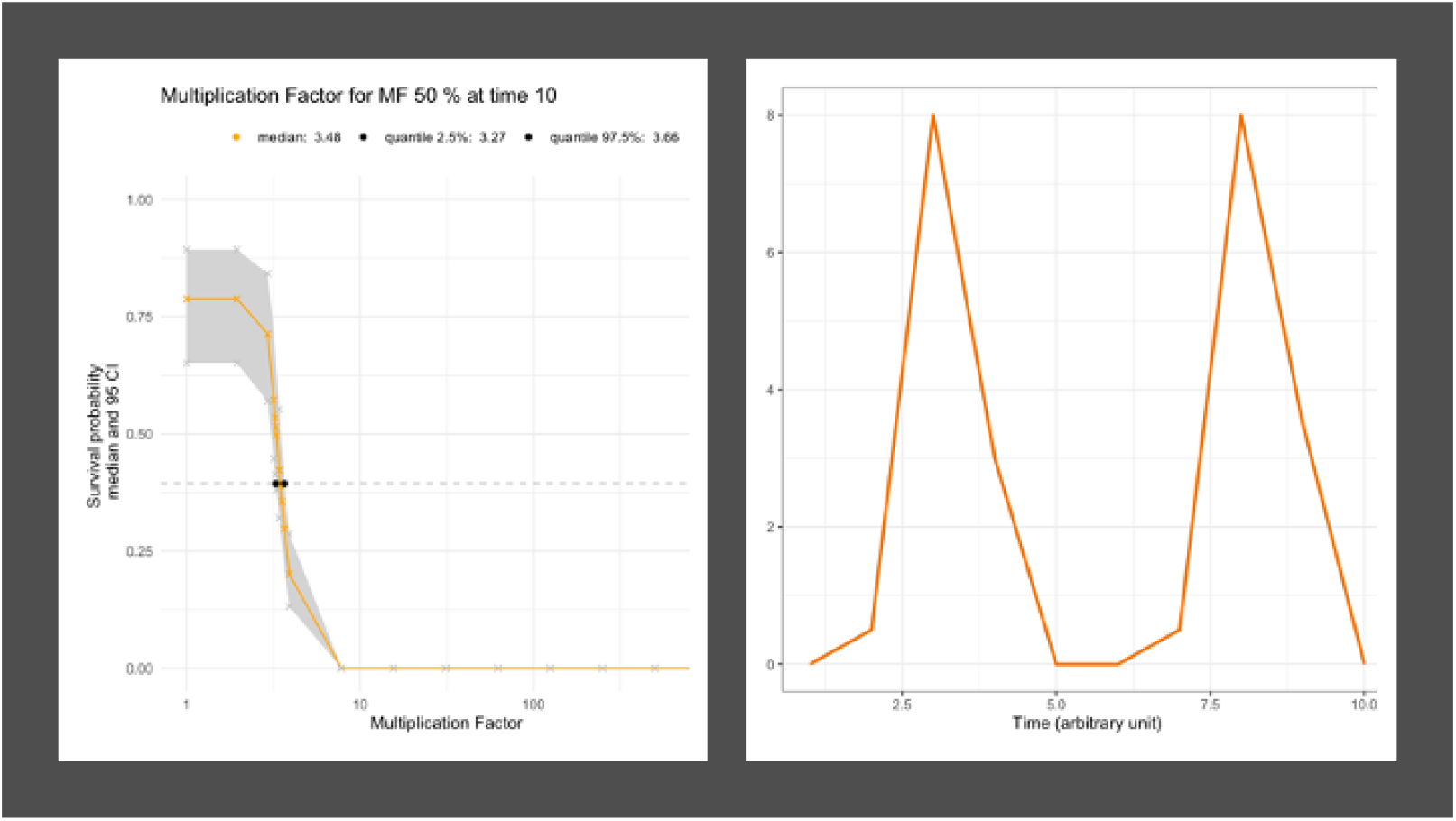
(left) Predicted survival probability according to a range of multiplication factors (log-scale) at the end of a theoretical exposure profile (right).

#### Predict survival probability under any exposure profile

Finally, it may be useful to predict the survival probability under any exposure profile (time-variable or not), for example when designing new experiments or to better understand what happens in field. Below are some examples from which you can inspire to perform your own simulations.

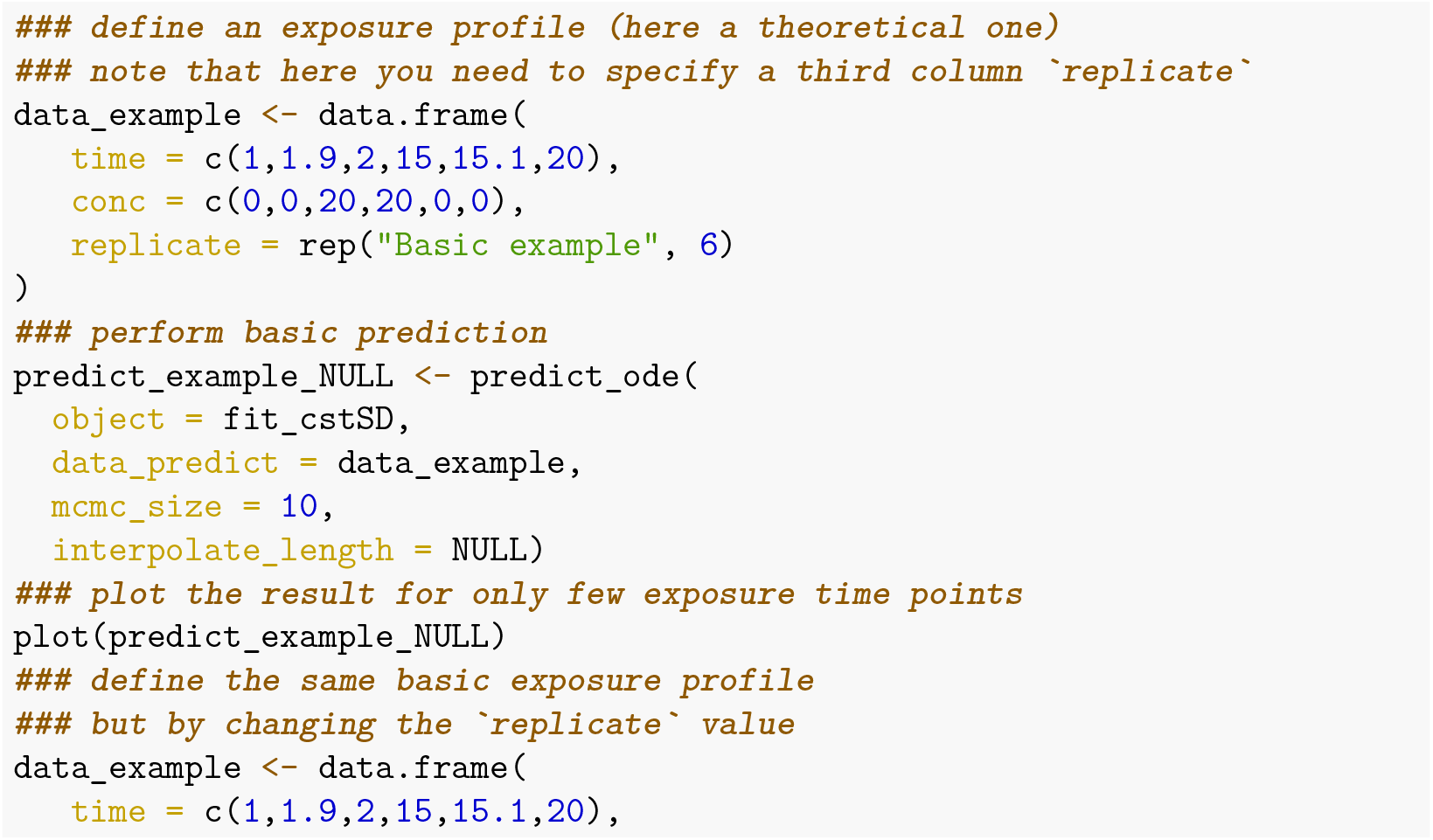

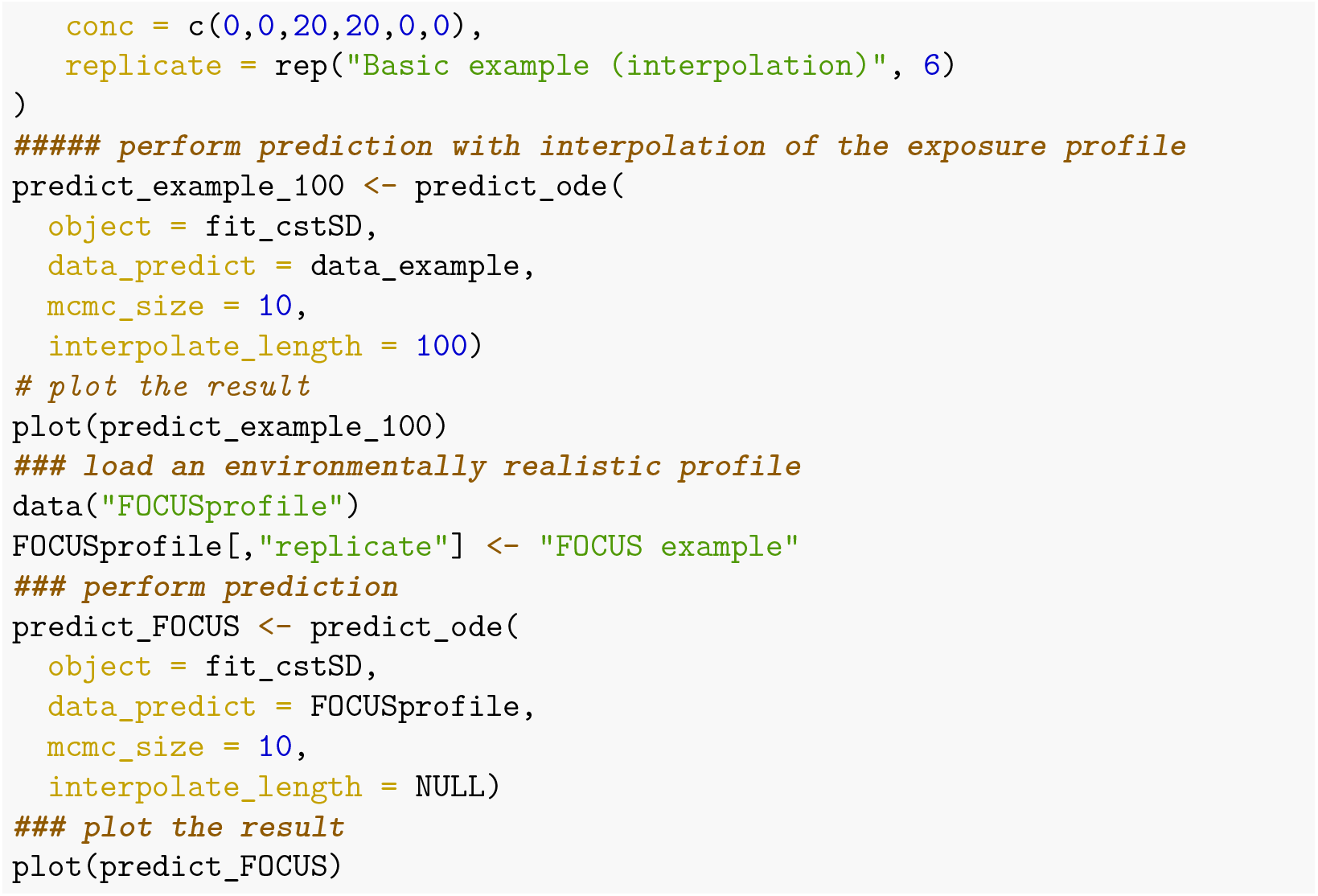

**Figure 6:**
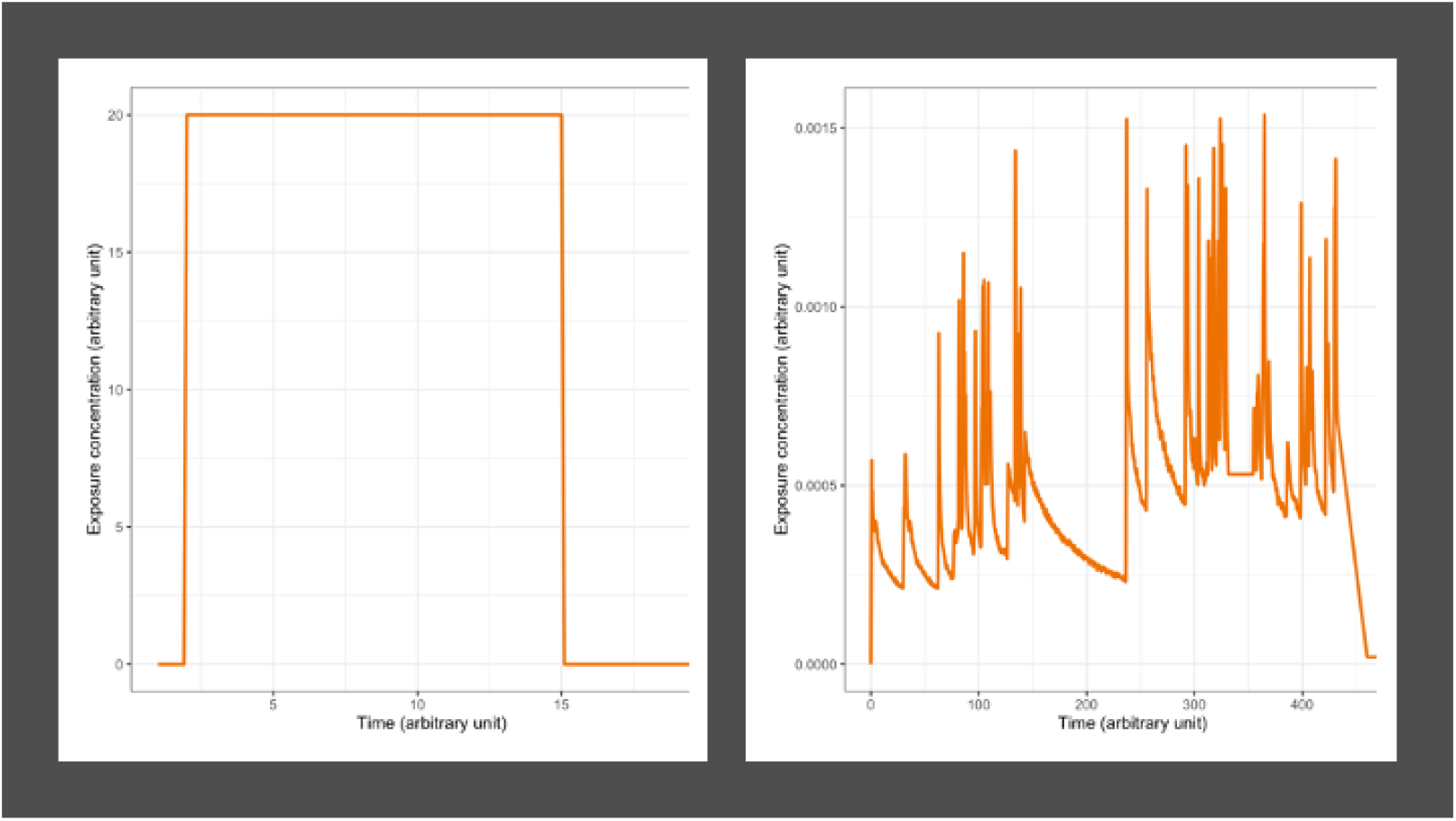
(left) Basic exposure profile; (right) Environmentally realistic exposure profile.

## Research using morse

Package `morse` was recently used to evaluate the added-value of using TKTD models in comparison with classical dose-response models, based on a case study with the snail *Limnaea stagnalis* when exposed to increasing concentrations of cadmium (Baudrot, Preux, Ducrot, Pavé, & Charles, 2018). Also based on `morse`, we proposed some recommendations to address TKTD assessment using uncertainties in environmental risk models (Baudrot & Charles, 2019).

